# Immune selection suppresses the emergence of drug resistance in malaria parasites but facilitates its spread

**DOI:** 10.1101/2020.12.15.422843

**Authors:** Alexander O. B. Whitlock, Jonathan J. Juliano, Nicole Mideo

## Abstract

Although drug resistance in *Plasmodium falciparum* typically evolves in regions of low transmission, resistance spreads readily following introduction to regions with a heavier disease burden. This suggests that the origin and the spread of resistance are governed by different processes, and that high transmission intensity specifically impedes the origin. Factors associated with high transmission, such as highly immune hosts and competition within genetically diverse infections, are associated with suppression of resistant lineages within hosts. However, interactions between these factors have rarely been investigated and the specific relationship between adaptive immunity and selection for resistance has not been explored. Here, we developed a multiscale, agent-based model of *Plasmodium* parasites, hosts, and vectors to examine how host and parasite dynamics shape the evolution of resistance in populations with different transmission intensities. We found that selection for antigenic novelty (“immune selection”) and within-host competition both suppressed the evolution of resistance in high transmission settings. We show that high levels of population immunity increased the strength of immune selection relative to selection for resistance. As a result, immune selection delayed the evolution of resistance in high transmission populations by allowing novel, sensitive lineages to remain in circulation at the expense of common, resistant lineages. In contrast, in low transmission populations, we observed that common, resistant strains were able to sweep to high population prevalence without interference. Additionally, we found that the relationship between immune selection and resistance changed when resistance was widespread in the population. Once resistance was common enough to be found on many antigenic backgrounds, immune selection stably maintained resistance in the population because resistance was able to proliferate, even in untreated hosts, when it was linked to a novel epitope. The results of our simulations demonstrate that immune selection plays a major role in observed dynamics of resistance evolution.

**Author summary:** Drug resistance in the malaria parasite, *Plasmodium falciparum*, presents an ongoing public health challenge, but aspects of its evolution are poorly understood. Although antimalarial resistance is common worldwide, it can typically be traced to just a handful of origins. Counterintuitively, although Sub Saharan Africa bears 90% of the global malaria burden, resistance typically originates in regions where transmission intensity is low. In high transmission regions, infections are genetically diverse, and hosts have significant standing adaptive immunity, both of which are known to suppress frequency of resistance within infections. However, interactions between immune-driven selection, transmission intensity, and resistance have not been investigated. Using a multiscale, agent-based model, we found that high transmission intensity slowed the evolution of resistance in two ways. First, it intensified competition, allowing sensitive lineages to suppress resistant lineages. Second, high population immunity selected for antigenic novelty, which interfered with selection for resistance. However, once resistance was at a high frequency, immune selection maintained it in the population at a high prevalence. Our findings provide a novel explanation for observations about the origin of resistance and demonstrate that adaptive immunity is a critical component of selection.

## Introduction

Although widespread drug resistance in *Plasmodium falciparum* malaria has been an ongoing public health challenge for decades, several observations about the evolution of resistance seem counterintuitive. For one, despite hundreds of millions of cases of malaria per year [1], resistance mutations arise surprisingly rarely. Resistance mutations occur readily in the lab [2], and the parasite population size and mutation rate ensure that every infection should contain multiple resistance mutations [3]. However, widely circulating resistant lineages can typically be traced to just a handful of origins [4–9]. Furthermore, the points of origin are not, geographically, where one might expect. Sub-Saharan Africa suffers 90% of the global burden of malaria [1], but antimalarial resistance rarely originates there. Instead, resistant lineages typically originate in regions with relatively low transmission intensity, such as South America or Southeast Asia. Once introduced to Africa, however, these resistant lineages spread readily. For example, resistance to chloroquine originated five times in South America and Southeast Asia [10] but only spread through Africa once introduced from Asia in 1978 [8]. Together, these observations suggest that there is a dynamic that suppresses the spread of resistance immediately after origin and, further, that this dynamic is intensified in high transmission settings. Unfortunately, the spread of an established resistance mutation is demonstrably less constrained. One by one, P. *falciparum* has evolved resistance to every drug introduced. The current front-line treatment, artemisinin combination therapy (ACT), is still generally effective in most countries, but resistance is rising: treatment failure rates exceed 50% in most of Southeast Asia [11]. With millions of lives at stake, fully understanding the evolution of resistance is urgent.

Transmission intensity shapes both the genetic make-up of the circulating parasite population and the within-host environment those parasites experience. Most malaria infections are genetically diverse [12–14], and this diversity increases in high transmission regions where hosts experience frequent, overlapping infections [15–18]. In a multiply-infected host, different lineages compete for the same resources while subject to both specific and nonspecific immune regulation. When drug-resistant and drug-sensitive parasites are present in the same host, sensitive lineages may suppress the growth and transmission of resistant lineages in a phenomenon known as competitive suppression [19–23]. Frequent infections also result in high levels of adaptive immunity. Adaptive immunity is not sterilizing; instead, infections in a highly immune host remain asymptomatic [24, 25]. For example, individuals in holoendemic regions of Africa receive upwards of 500 infectious bites per year [26] and symptomatic infections become increasingly rare after age five [25]. Conversely, in regions with low transmission intensity, hosts may never develop broad immunity. Within hosts, high levels of adaptive immunity are associated with increased clearance of resistant parasites [27]. While the influence of transmission intensity on diversity, competition, and immunity are relatively well-understood, the interactions between these forces may shape the evolution of resistance in complex ways.

Competitive suppression of drug resistant parasites by sensitive ones is typically considered a consequence of a fitness cost of resistance. Estimates of the cost of resistance have been based on observational epidemiological data as well as direct competition assays *in vitro* [28–31]. Measuring the intrinsic cost of any resistance mutation may be complicated by the presence of compensatory mutations [32–34]. Furthermore, the consequence of any potential fitness cost on selection will depend on additional factors, including the complexity of infection, host immunity, and treatment rate [35]. For example, high levels of host immunity can suppress resistant lineages even when resistance is cost-free. In high transmission settings, low-frequency resistant mutants will often arrive in a currently infected host whose active immune responses are likely to lead to the indiscriminate loss of rare variants [36]. Even in naive hosts, numerical dominance of a sensitive strain can lead to the loss of a resistant strain as density-dependent immune regulation of the entire parasite population begins [37]. The power of host immunity to limit the within-host parasite population size also decreases the likelihood that a resistance mutation occurs in the first place [38].

These ecological constraints occur independently of selection on resistant variants, but high levels of population immunity alter the adaptive landscape as well. Due to its strain-specificity, adaptive immunity generates selective pressure which favours the proliferation of lineages with novel epitopes in a partially immune host. All else equal, as the number of exposures increases for a particular host, the proportion of circulating strains that are able to infect that host decreases, as does the antigenic diversity of any infection that occurs [39]. As a result, in highly immune host populations, parasites are subject to an additional dimension of selection in the form of selection for antigenic novelty.

While mathematical and computational models have been extraordinarily useful for investigating the processes driving patterns of resistance evolution in malaria parasites [19,21,36,37,40–47], as far as we are aware, none have explored the potential for interactions between selection for antigenic novelty and selection for drug resistance. Additionally, many have necessarily made simplifying assumptions, such as simulating homogeneous host populations, introducing mutations at a relatively high frequency, or using deterministic models of evolutionary dynamics. However, host heterogeneity, such as an individual’s exposure history, will affect the environment of a new mutation [40], a rare mutation will be subject to different selective pressure than one at higher frequency, and *Plasmodium* life history is characterized by severe bottlenecks alternating with massive population expansion, intensifying the contribution of both selection and drift [43, 46, 48, 49]. As a result, it is difficult to draw conclusions about the origin of a resistance mutation using these prior approaches.

Here we develop a multi-scale, agent-based, stochastic model of *Plasmodium* infection, transmission, and evolution in a host (and vector) population. We explicitly track parasites and their genomes; hosts, their immune systems, and their red blood cell resources; mosquito vectors; and the interactions between all three organisms. Importantly, resistance mutations are distinct from antigenic background, allowing selection to work independently on both. While our model makes some explicit assumptions (e.g., host and vector population sizes, the number of antigenically-unique parasite strains), evolutionary drivers like transmission intensity, multiplicity of infection, genetic diversity in the parasite population, and characteristics of host immunity are emergent properties. We demonstrate that selection for antigenic novelty and within-host competition interact to delay resistance evolution. In high transmission settings, immune selection antagonizes the early spread of resistance by maintaining sensitive lineages in the populations. However, once resistance is common, immune selection promotes its maintenance regardless of transmission intensity. Additionally, our model reveals that antigenic diversity and novelty are necessary for resistance to reach a high prevalence. Our results provide a mechanism to explain the observed pattern of resistance evolution, in which high transmission intensity impedes the origin, but not the spread, of resistance.

## Results

### Endemicity and infection diversity increase with EIR and number of antigenic strains

To briefly summarize the simulations, each vector, parasite, and host was individually modelled. Each parasite had a discrete, three locus genome. Two of the loci were associated with both resistance and growth rate, such that a mutation at either or both produced full drug resistance as well as a cost in the form of a growth rate reduction. The allele at the third locus determined interactions with the host immune responses. We refer to all parasites that share an allele here as a “strain”, which is equivalent to a serotype. Mutations at each locus were random and recurrent. At the time of population initiation, every parasite was genetically identical. Over time, genetic diversity evolved through the forces of mutation, selection, and drift. The maximum number of possible strain variants was capped at either one strain, representing no antigenic diversity, or 30 strains. Because strain identity evolved independently of resistance, a single strain may contain both sensitive and resistant lineages. Parasites reproduce asexually within the host and undergo sexual recombination within the vector. During an infection, hosts developed strain-specific immunity, which shaped the density and duration of current and future infections (S1 Fig, S2 Fig). Populations were founded with two different vector population sizes, producing low and high transmission intensity, and simulations were allowed to run for 2000 time steps, by which point genetic diversity in the parasite population and measures of endemicity in the host population had reached an equilibrium state.

Differences in transmission intensity in our model produced epidemiological patterns which reflected those observed in real populations. Using untreated, replicate populations at equilibrium, we measured aspects of endemicity and population dynamics. Increasing the number of vectors or strains increased transmission intensity. With 300 vectors and one strain, each host received an average of nine bites per year from infectious mosquitoes, akin to the entomological inoculation rate, or EIR. With higher numbers of vectors and strains—1200 and 30, respectively—mean EIR was over 200 (Fig 1C). EIR was lower with one strain because, without antigenic novelty, transmission was rare from any infection after the primary infection (S3 Fig), due to suppression of the parasite population by host immunity.

**Fig 1.**
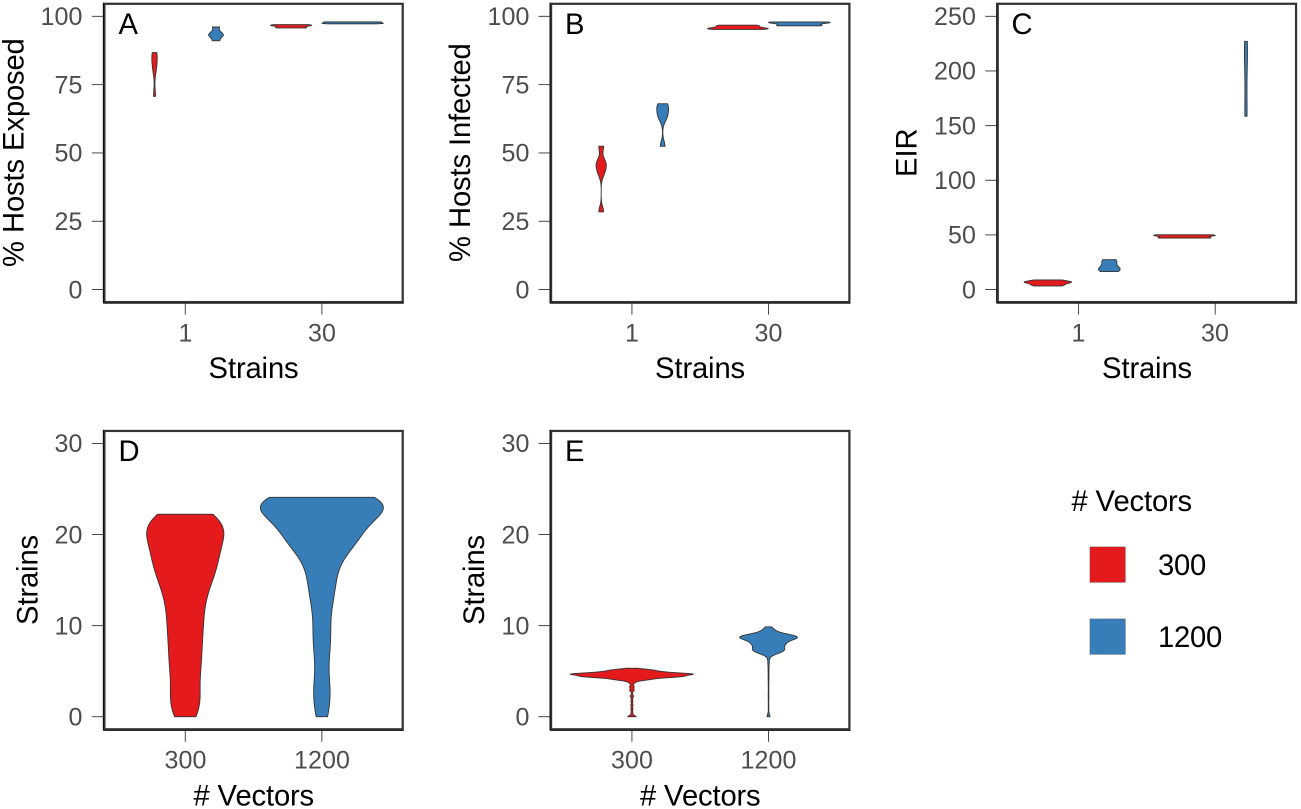
Baseline values in equilibrium populations. Ten populations for each condition were allowed to evolve to equilibrium, and mean values were measured over 100 time steps. (A) Percentage of hosts exposed (B) Percentage of hosts with active infections (C) Annual entomological inoculation rate (EIR) (D) Average number of strains to which hosts had been exposed (E) Complexity of infection, or average number of strains per infection.

At equilibrium, the hosts with active infections hosts and the proportion of hosts which had been exposed increased with number of strains and vectors (Fig 1A-B). With 30 strains, almost all hosts had been exposed and most were actively infected throughout the sampling window, indicating that hosts experienced repeated, overlapping infections, beginning early in life. With one strain, the proportion of infected hosts was less than the proportion of exposed hosts, indicating that hosts recovered from infections before potentially being infected again (Fig 1A-B). In accord with empirical measurements, high transmission intensity produced more complex infections (Fig 1E) and hosts were exposed to more strains earlier in life (S4 Fig, S5 Fig).

Because adaptive immunity is strain-specific, looking only at the fraction of hosts exposed to infection misses subtleties of the immune (and therefore fitness) landscape experienced by parasites. For example, there were only modest differences in the proportion of hosts exposed between the one strain, 1200 vector condition and the 30 strain conditions. However, the proportion of circulating strains to which those hosts were exposed varied substantially. In the former case, there was only one strain, so no antigenic novelty remained; when there were 30 possible strains, hosts on average were exposed to half and two-thirds of those strains in the low (300) and high (1200) vector conditions, respectively (Fig 1D). As a result, despite similar overall levels of exposure in the host population, differences in strain-specific exposures gave rise to different levels of population standing immunity, which we define as the likelihood that an arbitrary host has immunity to an arbitrary parasite. Operationally, we defined this metric as the average percent activation of immunity against a particular strain across hosts. Thus, the conditions could be ranked from highest population immunity to lowest as: one strain and 1200 vectors, one strain and 300 vectors, 30 strains and 1200 vectors, and 30 strains and 300 vectors.

Increasing the vector number changed the patterns of strain-specific immunity in the host population over time (Fig 2). In populations with 300 vectors, average immunity to individual strains fluctuated over time, frequently to the point that immunity to a particular strain was almost completely lost in the population. With 1200 vectors, mean immunity was higher, more stable, and more evenly distributed across strains, suggesting that frequent exposure maintained strain diversity. High levels of population immunity kept strains circulating at intermediate frequencies by strengthening selection for rare strains and against common strains. This form of balancing selection is consistent with field observations of antigenic diversity [50–52].

**Fig 2.**
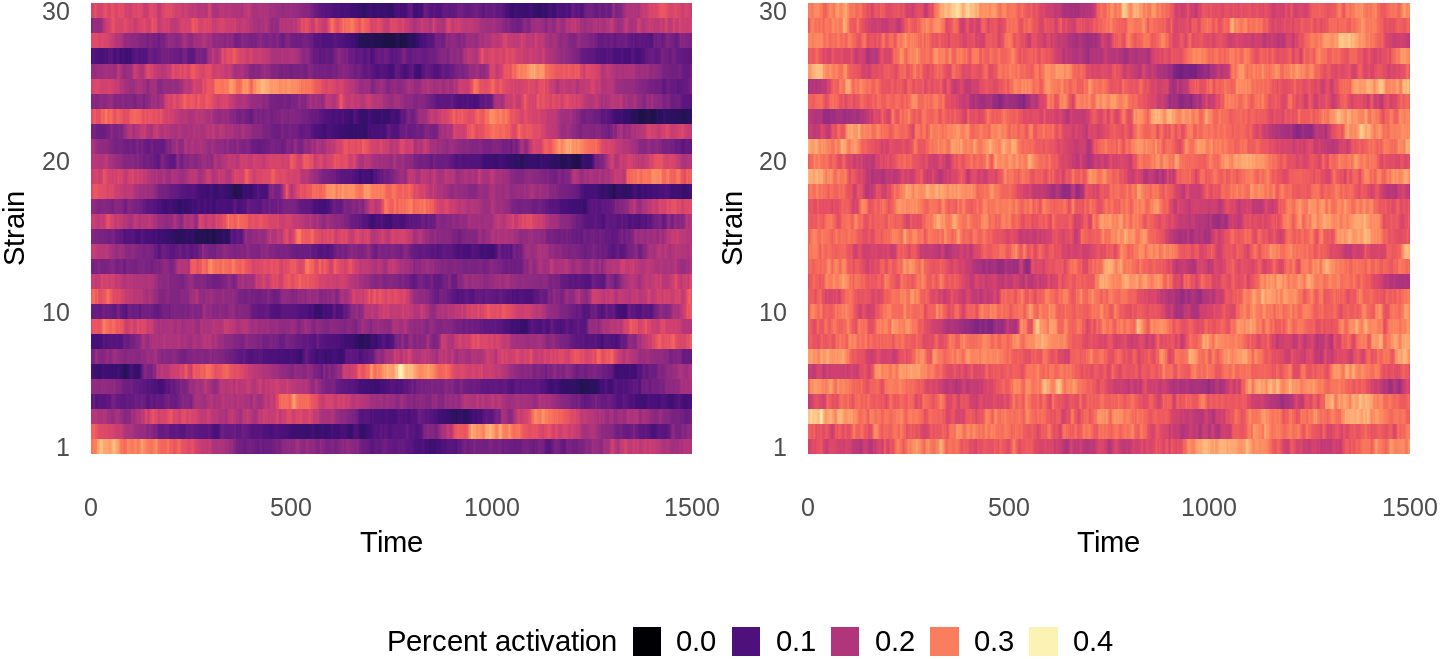
Population-wide mean strain-specific immunity over time in untreated equilibrium populations. Immunity to each strain is averaged over all hosts in the population at each time point. Left: 300 vectors. Right: 1200 vectors. Data shows one representative simulation for each condition.

### Low population immunity facilitated the spread of resistance

After populations reached equilibrium, we introduced drug treatment at a set rate, such that at any given time, 30% of hosts with symptomatic infections were treated. An infection was considered to be symptomatic if parasites exceeded a threshold abundance within the host. Population endemicity dropped immediately after treatment introduction, but rebounded over time as resistance evolved and spread. We allowed evolution to proceed until the prevalence of treatment-resistant infections in the host population reached an equilibrium.

We found that strain dynamics were the most important factor modulating the evolution of resistance, but there were additional, complex interactions driven by the number of vectors and feedback from the changing prevalence of resistance. Resistance readily evolved and spread to a low prevalence among infected hosts by 200 time steps in all conditions (Fig 3A). The 30 strain and 300 vector condition reached *T*_fail_, the time at which 10% of infections were composed of at least 10% resistant parasites, most quickly (*p* < .001 and *A* = 0.1864 (large effect size) for the most similar pairwise comparison), but the decrease was relatively small, and there was no significant difference between the other conditions (*p* = .541 for the least similar pairwise comparison). The similarity between conditions demonstrates that later results were not due to differences between mutation supply or a lack of selection for resistance.

**Fig 3.**
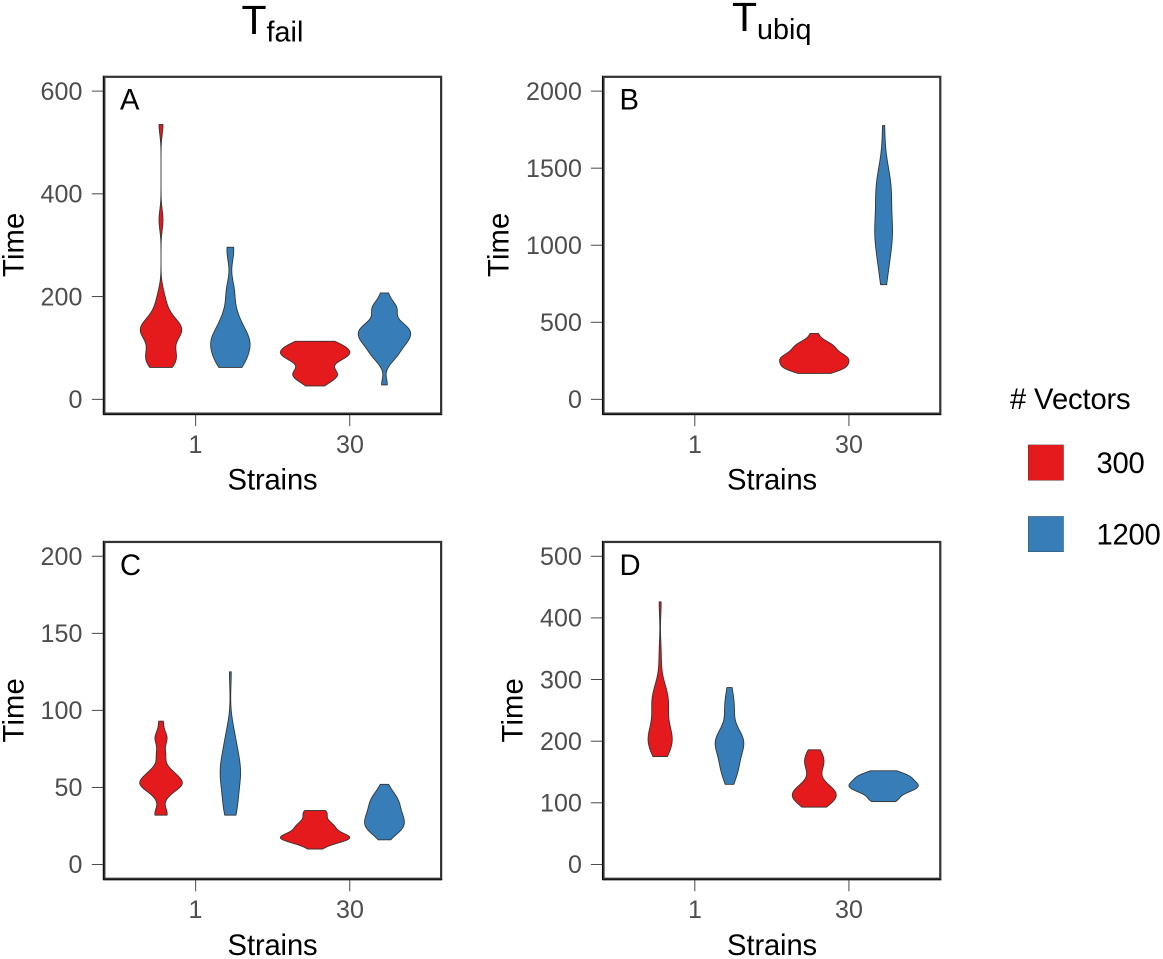
Time to resistance. Mean time to 10% prevalence of treatment failure (*T*_faii_) and 75% prevalence of treatment failure (*T*_ubiq_) was measured over 25 replicate populations. Top row: Costly resistance. A: *T*_fail_. B: *T*_ubiq_. Bottom row: Costless resistance. C: *T*_fail_. D: *T*_ubiq_. Note different Y axes.

In simulations with one strain, populations never reached *T*_ubiq_ (Fig 3B), defined as the time at which 75% of infections were composed of at least 10% resistant parasites. Equilibrium prevalence of treatment-resistant infections was relatively low, with large fluctuations over time (Fig 4). The prevalence of resistance was lower with 1200 vectors because resistant lineages could only reach densities sufficient for transmission in naive hosts, and there were fewer naive hosts in high transmission populations (p < .001, A = 0.122 (large effect size)). Within hosts, mixed infections—those harboring both sensitive and resistant parasites—were rare (Fig 5A and C); in most infections, the parasite population was either almost entirely sensitive or almost entirely resistant. This is because the relative fitness of a resistant lineage in an infection was entirely dependent on the host’s treatment status. Sensitive lineages were rapidly eliminated in treated hosts, but in an untreated host, sensitive lineages outcompeted resistant lineages. As a result, sensitive lineages continued to proliferate in the population.

**Fig 4.**
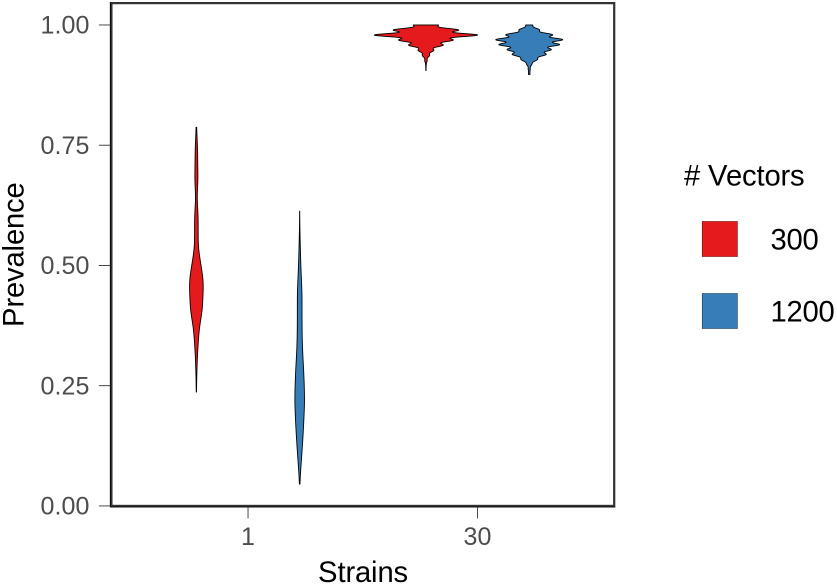
Equilibrium resistance prevalence. After treated populations had reached equilibrium, the average prevalence of treatment failure (10% within-host frequency of resistance) was measured over 100 time steps in 25 replicate populations.

**Fig 5.**
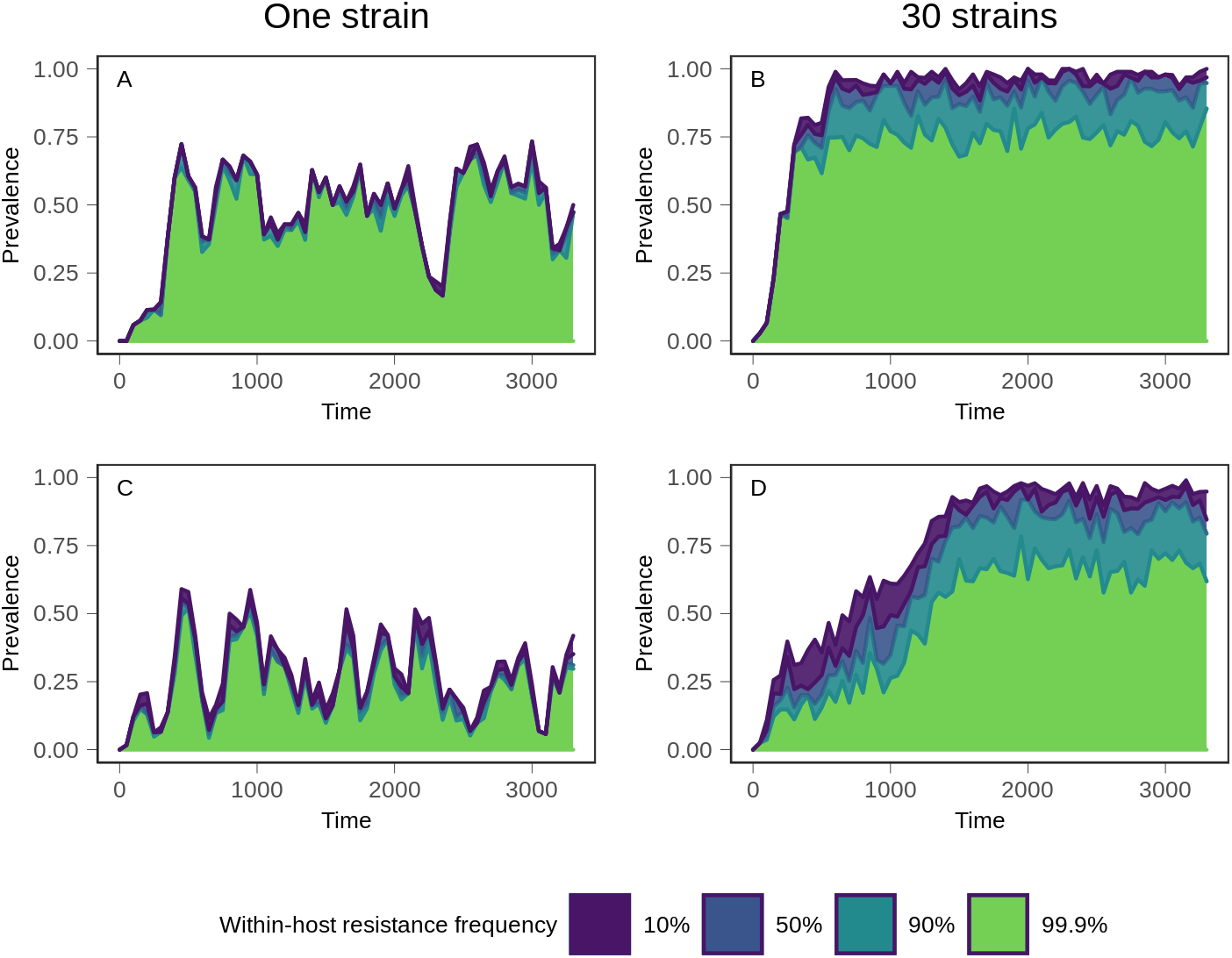
Evolutionary trajectory of resistance. Following introduction of treatment at time 0, the prevalence of within-host frequencies of resistance was monitored over time. The top line indicates the prevalence of all treatment-resistant infections, defined as infections consisting of at least 10% resistant parasites. Moving down from the top, the prevalence of infections consisting of between 10% and 50% resistant parasites is shown in blue. Below that, the teal section shows the prevalence of infections in which the frequency of resistant parasites is between 50% and 90%. Finally, the bottom section in green shows the prevalence of infections in which resistance is at near fixation within the host, with a frequency exceeding 99.9%. Top row (A and B): 300 vectors. Bottom row (C and D): 1200 vectors. Left column (A and C): one strain. Right column (B and D): 30 strains. The minimal separation between lines in the one strain conditions demonstrates that mixed infections are rare, compared to 30 strain conditions. Data shows one representative simulation for each condition.

With 30 strains, treatment-resistant infections reached a high, stable prevalence that exceeded ubiquity in both vector conditions although, as in the one strain conditions, prevalence was lower with 1200 vectors (*p* < .001, *A* = 0.399 (small effect size)). Higher levels of exposure (i.e., more vectors) also slowed the spread of resistance (Fig 3B). *T*_ubiq_ was approximately 250 time steps with 300 vectors, compared to over 1000 time steps for 1200 vectors. Although increased strain number increased the prevalence of treatment-resistant infections in the population, it also suppressed the frequency of resistance within individual hosts. A substantial and stable proportion of infections contained both sensitive and resistant parasites at equilibrium (Fig 5B and D). With 1200 vectors, mixed infections were common before equilibrium as well, but they were rare with 300 vectors.

The differences can be understood in terms of an adaptive landscape, in which the parasite traits under selection were growth rate, resistance, and immune novelty. A resistance mutation arising on a novel strain in a treated host had multiple within-host advantages, and thus was under the strongest positive selection. In contrast, resistance arising in an untreated host on a strain to which that host had previously been exposed was likely to be quickly purged due to the combined effects of immunity and the cost of resistance. Other scenarios gave rise to different outcomes. Antigenic diversity allowed resistance to reach a high prevalence in the host population because resistance mutations could proliferate if they were linked to a novel strain, even in untreated hosts. In this manner, strain-specific immune selection stabilized resistance in the population by providing a countervailing force to the cost of resistance. As resistance rose in frequency in the parasite population, a higher proportion of strain mutations occurred in resistant lineages, maintaining a supply of antigenic diversity and preventing the loss of resistance due to competition or immune selection. The lower the strain-specific immunity among hosts, the higher the likelihood that resistance mutations originated in a host environment in which they were highly beneficial.

### Removing cost decreased the impact of population immunity on resistance evolution

The high complexity of infection that is characteristic of high transmission settings increases the likelihood that a host will be co-infected with both sensitive and resistant lineages, leading to the hypothesis that within-host competition suppresses the evolution of resistance. To test this hypothesis, we allowed populations to evolve as before, but removed the cost of resistance, eliminating competitive suppression due to growth rate.

As expected, the absence of cost accelerated the rate of resistance evolution (i.e., decreased *T*_fail_ and *T*_ubiq_) in all conditions, but the effect was most significant with one strain, where populations were able to evolve stable resistance and reach ubiquity (Fig 3D), which did not occur when there was a cost to resistance (recall Fig 3B). With only one strain, a higher vector number allowed resistance to reach ubiquity more rapidly (*p* = .005, *A* = 0.268 (medium effect size)), demonstrating that high transmission intensity dispersed mutations more rapidly throughout the population. Although we cannot rule out the possibility that high transmission suppressed resistance by exacerbating within-host ecological processes, such as prior residency or host immunity, the fact that high transmission was advantageous when resistance was cost-free indicated that within-host competition was the dominant suppressive force when resistance was costly.

Removing cost sped up the evolution of resistance in both 30 strain conditions, most significantly with 1200 vectors. This indicated that competitive suppression delayed the evolution of resistance, especially with high transmission. However, unlike in the one strain conditions, high transmission intensity did not accelerate resistance’s spread to ubiquity (*p* = .449) because immune selection dynamics counterbalanced the advantage conferred by increased transmission opportunities. In other words, immune selection alone slowed the spread of a beneficial mutation.

When resistance was costly, a resistance mutation could only proliferate in untreated hosts if it was genetically linked to a novel strain. Without cost, and without multiple strains, resistance mutations were only selectively neutral or beneficial, depending on the treatment status of the host. Therefore transmission rate was the major factor determining their rate of spread. With multiple strains, a lineage carrying a resistance mutation may still have be selected against within the host environment if host immunity recognized the linked strain, but selection against it would have been weaker than when resistance was costly. Similarly, a resistance mutation linked to a novel strain would have been under positive selection, even in untreated hosts. This hastened the time to high prevalence of resistance.

Patterns of within-host frequency were qualitatively similar to simulations in which resistance is costly (S7 Fig). With 30 strains, mixed infections were present at equilibrium, regardless of the number of vectors, as well as before equilibrium with high transmission. Mixed (sensitive and resistant) infections were rare with one strain. This supports the notion that the suppression of within-host frequency observed with costly resistance in 30 strain conditions can be attributed to immune selection, rather than solely to growth rate competition.

### Immune selection for novel strains antagonized selection for resistance

In order to understand how immune interactions drive the evolution and maintenance of resistance at the parasite population level, we monitored strain dynamics over time, including strain-specific immunity averaged across all hosts, the frequency of each strain within the entire parasite population, and the proportion each strain contributed to the total level of drug resistance in the parasite population.

Shortly after treatment was introduced, the number of circulating strains dropped precipitously (Fig 6), as did the host population infection rate and the total size of the parasite metapopulation (i.e., all parasites, summed across all infections). These losses were especially severe with 300 vectors simulations, in which strain diversity was lower prior to treatment introduction.

**Fig 6.**
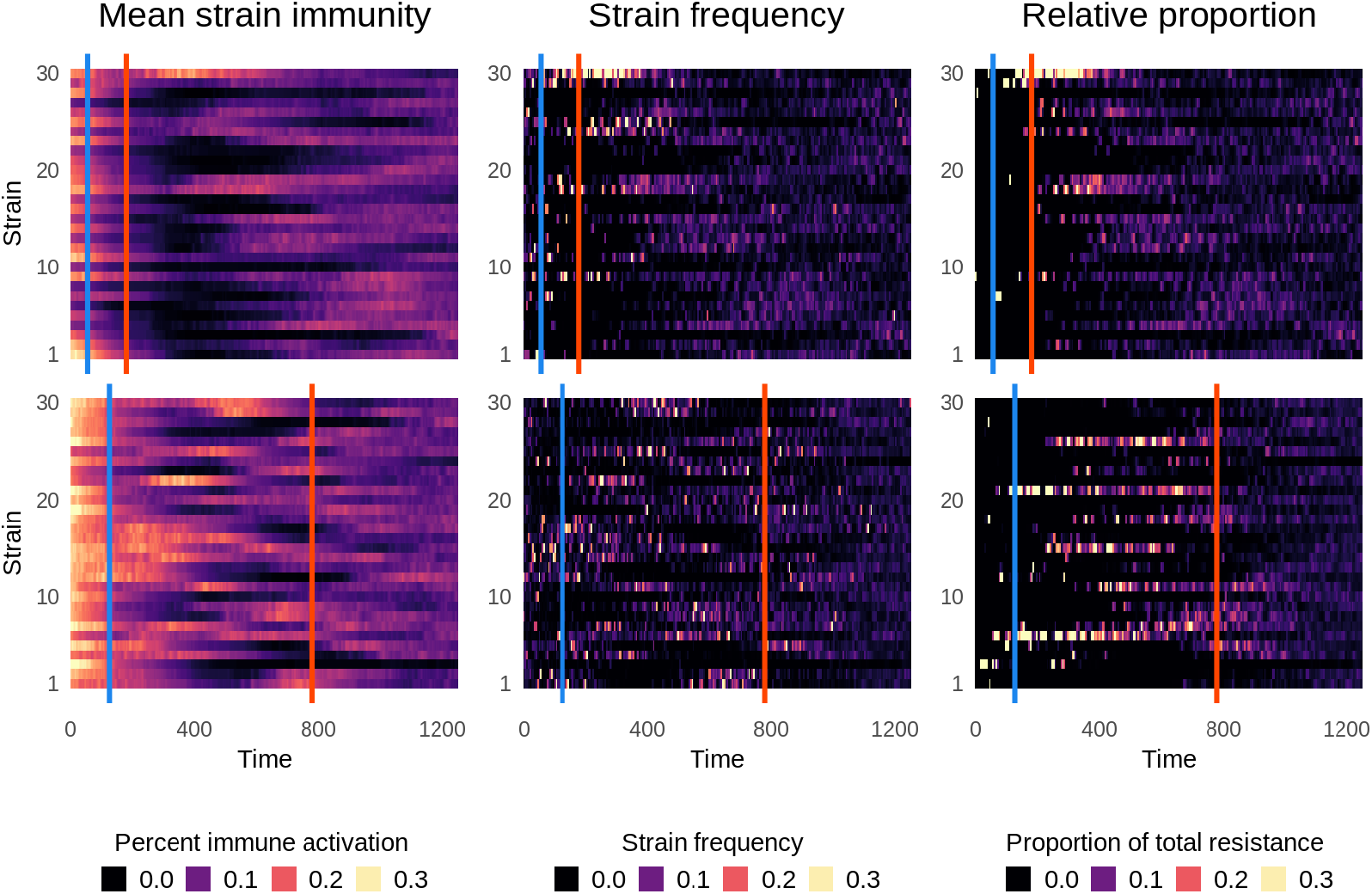
Strain-specific patterns of resistance. Following introduction of treatment at time 0, immunity and resistance were monitored within each strain over time. Top row: 300 vectors. Bottom row: 1200 vectors. Left column: Mean population immunity to each strain, as in Fig 2. Middle column: the frequency of each strain in the population. Right column: The relative contribution of each strain to the total amount of resistance in the population. The first line, in blue, denotes *T*_faii_, and the second line, in red, denotes *T*_ubiq_. Data shows one representative simulation.

With 300 vectors (Fig 6, top row), resistance rapidly evolved on the few strains which persisted after treatment introduction. In a phenomenon resembling a selective sweep, these resistant strains steadily increased in prevalence until resistance reached ubiquity. Most strain diversity was transient, but the frequency of the highly resistant strain 30 steadily increased (middle column), despite high levels of immunity against it. Strain 30 only declined after ubiquity, when resistance was found on many strains (Fig 6, right column, S8 Fig). This demonstrates that there was selection for immune novelty, but also that it failed to outweigh selection for resistance until resistance was ubiquitous.

With increasing vector number, more strains carried resistance mutations, producing more co-circulation and widespread standing adaptive immunity. Novelty in one host was less likely to translate to novelty in another host due to broader immune portfolios, so none of the strains were able to sweep to a high prevalence. Unlike with lower vector numbers, strain diversity remained high, even though most circulating strains were sensitive. Resistance only became ubiquitous once it was found on multiple strain backgrounds, indicating that balancing selection for strain diversity remained strong relative to selection for resistance (compare bottom row, center and right column in Fig 6). Additionally, changes in strain frequency tracked closely with changes in strain immunity, allowing rare sensitive strains to increase in prevalence. For example, at time 500, there was very little immunity to strains 1 and 2. Subsequently, they rose in frequency despite not carrying resistance mutations (S8 Fig). This demonstrates that, under high transmission intensity, immune selection outweighed selection for resistance.

The effect of vector number on pre-equilibrium strain diversity explains the difference in pre-equilibrium within-host frequency. With 300 vectors, only a few, highly resistant strains were circulating, and selection for resistance was much stronger than selection for novelty. This allowed quasi-fixation of resistance within hosts and promoted the rapid rise to ubiquity (Fig 5B). As the frequency of resistance rose, the strength of selection for resistance weakened. Immune selection only became strong enough, relative to selection for resistance, to suppress within-host resistance frequency after *T*_ubiq_. This was not the case with 1200 vectors, in which within-host frequency was suppressed throughout (Fig 5D). Immune selection was strong enough to maintain sensitive strains in the population, even when selection for resistance was at its strongest. While a higher transmission rate could disperse resistance more quickly, it also meant that sensitive lineages continued to be transmitted. The ongoing transmission of sensitive lineages delayed *T*_ubiq_.

At both transmission intensities, as resistant lineages reached high frequency, most strain mutations happened on a resistant background. The suppressive impact of immune selection lessened because resistance was no longer associated with a minority of strains. As a result, the availability of antigenic novelty maintained a high, stable prevalence of resistance.

## Discussion

Here we have demonstrated that there is a complex relationship between immune selection within hosts and the spread of resistance between hosts, and that the net relationship changes based on the prevalence of resistance in the host population. Immune selection was antagonistic during the initial spread of resistance. As the frequency of a rare resistant lineage increased, so did population immunity against that lineage, and therefore the strength of immune selection against it. This effect was strongest when transmission intensity was high and resulted in a delayed time to ubiquitous treatment failure. However, once resistance was common, immune selection maintained costly resistance at a high prevalence within the population, in part because linkage to a novel strain could counteract the cost of resistance. Although the antagonistic effect of immune selection was intensified by cost, immune selection suppressed within-host frequency of resistance even when resistance was not costly because sensitive lineages on a novel strain background could outcompete resistant lineages linked to common strains. These results expand on other studies that found that within-host ecological processes were sufficient to suppress resistance [36, 37]. A significant difference in this investigation is that recurrent strain mutation meant that immune selection alone was likely to produce an antigenically-diverse infection, and that a resistance mutation could be found on any strain. This increased the complexity of within-host competition and introduced negative frequency dependent selection on strain background. In that sense, competitive suppression early in an infection may have actually benefited the later expansion of a resistant lineage if a resistance mutation was linked to a rare strain. This mechanism did not exist in prior studies and may explain why, in our simulations, high transmission only delayed resistance when resistance was costly.

Immune selection provides a plausible explanation for the observation that resistance does not usually originate in high-transmission regions, but spreads rapidly once introduced. Asian *Plasmodium spp*. lineages have different common epitopes than those in Africa [53], and it is likely that Asian antigens are less cross-reactive with native African antigens. This would create positive immune selection for a newly-introduced strain and promote its rapid spread, even if resistance is costly, allowing it to reach a high, stable prevalence before immunity against that strain was widespread in the host population.

With rates of treatment failure rising and few alternatives available, a further understanding of immune selection could provide another tool in the fight to preserve the efficacy of existing antimalarials. This more critical than ever now that artemisinin resistance has been detected in Sub-Saharan Africa [54, 55]. One strategy is drug cycling, in which drugs are withdrawn from use as the rate of treatment failure rises in a population, and then reintroduced once resistance wanes [56]. Other investigations have supported the simultaneous use of multiple first-line therapies [44, 57]. Quantifying the relationship between population immunity, resistance prevalence, and strength of selection for resistance could be used to identify frequencies of resistance which act as tipping points, in order to most effectively determine when to use individual drugs and to identify optimal treatment combinations. Immune selection may also produce an unforeseen benefit of vaccination. To date, attempts to make a widely effective vaccine against *P. falciparum* have been challenged by its antigenic diversity and its extraordinary ability to evade the immune system [58,59]. The most effective vaccine to date, RTS,S/AS01, is only able to prevent approximately 30% of infections [60]. However, while the partial immunity produced by an imperfect vaccine may not totally prevent infection, it might alter immune selection dynamics, increasing the strength of selection for antigenic novelty and potentially interfering with selection for resistance.

These findings demonstrate that immune selection must be considered in any investigation of evolutionary processes in *Plasmodium spp*. There are many other factors we did not consider here. Due to limitations of computational feasibility and the complexity of the task, this work was constrained to a small region of the parameter space, including a steep cost of resistance and a fixed cross reactivity that assumes equal relatedness between strains. Wide ranges of values have been measured for many of these traits, and further, these values may change over time [32]. Many avenues for future investigation remain. For example, full resistance was produced by a point mutation, but resistance to some treatments, including ACT and sulfadoxine-pyrimethamine, is multigenic, while other resistant phenotypes are associated with a host of background mutations [61]. Although we found no effect from recombination in this model (S6 Fig), the level of linkage disequilibrium and epistatic interactions could influence the evolution of multilocus resistance [40], as would the presence of a tolerant intermediate [40, 43] or compensatory mutations [34]. Additionally, vector dynamics were highly simplified. *Plasmodium spp*. have many genes that are under selection within the mosquito [62, 63], meaning that the malaria parasite is subject to selection in multiple environments over the course of its life cycle. This complicates predictions about the spread of resistance based on within-host dynamics. Finally, the environment was static, but malaria is often seasonal in low transmission regions, which has implications for selection and complexity of infection [16]. This model is intended to be versatile and adaptable, and could be expanded to include any of these factors.

## Model and methods

### Overview

The purpose of our model is to explore the role and relative importance of selection driven by host immune responses and by drug treatment on the evolution of malaria parasites. We seek to further understand the observed pattern of drug resistance, in which it rarely arises in high transmission settings, but spreading easily once introduced from elsewhere [8]. Our model is individual-based at the levels of parasites, vectors, and hosts. Parasites are nested within hosts or vectors, parasite genotypes are explicitly tracked, and the state of the within-host environment (e.g., immunity and red blood cells) is defined at the individual host level. Simulations are conducted over discrete time steps, equivalent to days. Detailed methods and parameters can be found in the supplement; here we describe the building blocks (i.e., agents) of the model, the processes that the model captures, and the simulations that form the basis of our analysis.

### Individuals, state variables, and assumptions

#### Mosquitoes

Individual mosquito vectors are assumed to live for 20 days. Upon death, they are immediately replaced by a new mosquito of age zero. There is no background mortality nor age-related mortality increase. Malaria infection is assumed to have no impact on mosquito survival. Thus, the mosquito population size remains constant over the course of a given simulation, representing a carrying capacity in a given environment.

In the absence of infection, mosquitoes feed every three days [64], on days at which their age is divisible by three, so that population feeding is asynchronous. If the mosquito becomes infected, a one day feeding delay is added to represent the decreased feeding behaviour that has been observed in pre-infectious mosquitoes infected with *Plasmodium spp*. [65–67]. We assume that mosquitoes ingest two microliters of blood from a single host at each feeding.

#### Hosts

In our simulations, hosts live for a maximum of 500 time steps. While this is short, given our interpretation of time steps as days, we do this for two reasons. First, a high rate of transmission was required to prevent eradication when drug therapy was introduced to the population. As a result, endemicity in the low transmission conditions substantially exceeded the endemicity of the natural resistance hotspots in Southeast Asia and South America. At older host ages, just as in holoendemic regions, constant, asymptomatic infection became the rule under all conditions. Because the population size was limited by computational constraints, increasing the host lifespan caused the population to be saturated with infection. Excluding older hosts increased the difference between the high and low transmission conditions because the rate of exposure was slower in low transmission conditions. This allowed us to focus on the part of the lifespan where the populations diverge the most.

Hosts die from random background mortality at a fixed rate. Hosts can also die from malaria infection if their within-host count of uninfected red blood cells falls below a minimum threshold or if the percent of cells infected (parasitemia) exceeds a maximum threshold. When a host dies due to background mortality, infection, or reaching the maximum age, they are are immediately replaced by a new host of age zero. Thus, the host population size, *N*, remains constant over the course of the simulation.

We prioritized parameter choices that created realistic variation in infection outcomes because infection duration, immunity levels, and asymptomatic transmission all had potential to affect the within-host environment and the dynamics of selection. In human populations, the duration of infections is highly variable and difficult to measure, and both acute and chronic infections are common [68–72]. Variation in host response to infection was generated in two ways. First, each host had a randomly selected hyperparasitemia mortality threshold drawn from a Poisson distribution with mean of 11%, a value based on the WHO threshold for severe *P. falciparum* malaria [73]. Second, each host had a randomly selected rate of growth of the adaptive immune response. These factors produced a range of outcomes and course of disease that is comparable to empirical observations.

Within a naive host, our default parameters give rise to peak parasitemia between day 12-14 post-infection. Approximately 25% of hosts die at this time during a primary infection and an additional 25% of infections are cleared (S1 Fig). Early clearance is highly time-sensitive and occurs when adaptive immunity increases quickly enough to produce a substantial effect while innate immunity is still at peak (S2 Fig). The rest of the infections last at least 150 days and are characterized by dampened oscillations produced by antigenic escape [72]. Mortality is rare during later stages of infection. Infections are asymptomatic during most of that time, though occasionally oscillations will cause transient symptoms. Asymptomatic infections are still transmissible. Subsequent infections are typically low density and truncated. Reinfections with a previously exposed strain usually remain subpatent and are unable to produce gametocytes, but a novel strain may produce a symptomatic, transmissible infection (S3 Fig).

#### Parasites

Individual parasites are tracked within hosts and within vectors. Each parasite has a unique genotype consisting of three unlinked loci, two of which encode resistance (and therefore determine growth rate). The allele at the third locus determines interactions with the host immune responses. We refer to all parasites that share an allele here as a “strain”. Strains are equivalent to serotypes, and all members of the same strain have identical epitopes and elicit the same immune response. The genome uses the standard nucleotides A, G, T, and C. Each base is subject to mutation at a given rate at each replication cycle. The genomic mutation rate—applying to the loci determining resistance and growth rate—is 0.0005 per residue per replication. This value makes it likely that a small number of resistant mutants will be generated in each infection, especially during primary infections, in accord with expectations [3]. We assume a finite number of possible strains. At each replication cycle, there is a probability of mutation to a different random strain, chosen with equal probability. We assume the probability of antigenic change is small, 0.00001 per replication, allowing variation in growth rate/resistance to increase within a single antigenic strain as a consequence of selection driven by competition and/or drug treatment and increasing the likelihood that strain diversity (i.e., circulation of multiple antigenic lineages) is the product of selection for antigenic novelty.

A parasite genotype is drug resistant if either or both resistance-associated loci are mutated to a value of “C”. There is no epistasis between loci, so any genotype with at least one resistance mutation produces a fully resistant phenotype. The cost of resistance is modelled as an intrinsic growth rate reduction. In untreated naive hosts, an infection containing only resistant mutants will grow at 40% of the rate of an infection containing only sensitive parasites. In an untreated mixed infection, expansion of a resistant lineage will additionally be limited by immunity elicited against the faster-growing sensitive population.

### Processes

#### Transmission to hosts and within-host infection dynamics

An infection in a host begins when an infectious mosquito takes a blood meal. We assume this occurs with a probability of one. The number of motile, transmissible parasites (sporozoites) injected into the host are drawn from a Poisson distribution with a mean of twelve, which ensures a blood-stage infection that leads to gradual build up of immunity within the host. In reality, these parasites travel to the liver and undergo several rounds of replication; in our model, sporozoites spend seven days dormant, during which time they are not recognized by the host immune system. At the end of this period, each sporozoite produces five merozoites—the red blood cell (RBC) infecting parasite stage.

We assume hosts have a maximum of 8.5 × 10^6^ RBCs and produce up to 3.7 × 10^5^ new RBCs per day. We choose these numbers for computational practicality and note that they essentially correspond to modeling dynamics in one *μ*L of blood in a mouse. In an uninfected host, RBCs are subject to background mortality. In an infected host, uninfected RBC are subject to bystander killing, in which infected cells increase the mortality rate of uninfected cells [74–76]. Infected RBCs have a higher background mortality rate due to fragility [77] and clearance by the spleen [73].

The number of erythrocytes, *a*, infected in each host at each time point is calculated as

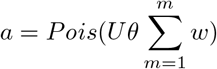

where *U* represents the number of uninfected erythrocytes per microliter, *w* represents the growth rate of an individual merozoite, *m*, and *θ* is a constant representing infection probability, which can be considered as the probability that a merozoite encounters and infects an RBC [36]. At each time step, *a* merozoites are chosen randomly with replacement to infect RBCs, but the probability that any individual merozoite will infect an RBC is weighted by its growth rate. The number of RBCs infected cannot exceed the summed growth rate of all merozoites. In other words, ten merozoites with a growth rate of 1 will infect no more than ten RBCs; ten merozoites with a growth rate of 0.5 will infect no more than five RBCs.

Once infecting an RBC, parasites take one of two developmental pathways. In 95% of infected cells, parasites replicate asexually, releasing ten progeny parasites one day after infection. In 5% of infected cells, parasites instead go on to produce gametocytes, the stage of the parasite that is transmissible to mosquitoes. While this value is constant and does not reflect predictions about parasite phenotypic plasticity [78, 79], it corresponds to overall values observed in mice [80]. We assume gametocytes take two days to mature; thus, the within-host life cycle roughly captures the biological details of *P. chabaudi* infection in mice. Again, these choices were for computational practicality. While altering the cell cycle duration of parasites or blood density of hosts may quantitatively alter the timing of evolutionary dynamics, we were concerned with the relative timing across different contexts.

Merozoites have an instantaneous background mortality drawn from Poisson distribution with a mean of 20%. Any merozoites which do not infect an erythrocyte are cleared in each time step. Gametocytes are removed via daily background mortality. We do not separately track different gametocyte sexes.

The host immune response is composed of both innate and adaptive responses. We model immunity phenomenologically in terms of killing rate. Both arms of immunity are dependent on the density of infected erythrocytes, and all blood-stage parasites are susceptible to their effects. Total immune activity against a given parasite strain is the sum of four processes: innate activation, adaptive activation, cross-reactive activation, and immune saturation (which reduces total activity). An overview of each is described below, with a full description in the supplement.

Innate immunity is strictly density dependent, peaking early and rapidly declining. Its growth and its effects are strain-independent. Adaptive immunity is specific to each antigenic strain. There is no compartmentalization, i.e., the immune response to a given strain is not decreased by co-infection with additional strains. At each time step in an infected host, adaptive immunity grows as a function of an individual host’s adaptive growth rate and infected erythrocyte density, and is decreased by antigenic escape (see below). Once a host is no longer exposed to a strain, adaptive immunity to that strain decays slowly, even if the host is infected by other strains.

While adaptive immunity grows and decays independently for each strain, strains are cross-reactive, and total adaptive immune activity against a given strain is increased by adaptive immunity to other strains. We assume that each strain is identically cross-reactive to the others.

Antigenic escape is conceptually based on PfEMP1 proteins encoded by var genes [81,82]. PfEMP1 variation is independent of serotype. The rate of antigenic escape shapes the duration and course of infections [83]. Early in an infection, there are many possible alternative configurations. Initially, adaptive immunity fluctuates with changes in density of infected erythrocytes. As the duration of exposure increases, the rate of escape slows as fewer novel variants remain and eventually halts once all possible configurations have been exhausted, resulting in sustained growth of adaptive immunity even with a low density of infected erythrocytes.

Exposure is measured independently for each strain, such that if strain two is introduced into a host with a long duration of exposure to strain one, the immune response against strain two will be bolstered by cross-reactivity against one, but the growth of the strain two-specific immune response will be slow initially because strain two retains full potential for antigenic escape.

Finally, the immune system’s capacity to clear parasites saturates at high parasite densities. There is a sigmoidal relationship between the efficacy of immune response and parasite density [84].

#### Transmission to vectors and within-vector infection dynamics

Mosquitoes can become infected with malaria when feeding on an infected host. The probability of infection, *p*, is a sigmoidal function of within-host gametocyte density, given as *L*, per microlitre. Following [29], we used the relationship determined in [85] with shape values based on data from [86]. The probability of infection is thus given as,

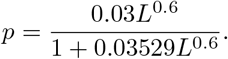

If the vector is infected, it will ingest 2*L* gametocytes, assuming a 2*μ*L bloodmeal size. The number of gametocytes is assumed to not be a limiting factor, so gametocytes which are ingested by a vector are not removed from the host’s pool of gametocytes. From the ingested gametocytes, 1% are randomly chosen to sexually reproduce with another gametocyte from the same bloodmeal, with free recombination between all three loci. Vector immunity dynamics are not explicitly modelled, but the other 99% of the ingested parasites are assumed to die before becoming oocysts. Remarkably, this is less severe than the average 2,754-fold decrease observed in experimental infections [87]. Ten days after the bloodmeal [62], each oocyst bursts to produce 12 sporozoites, each of which undergoes mutation independently with the same parameters as within-host mutation. Once sporozoites are produced, the mosquito is infectious.

#### Drug treatment

By default, 30% of symptomatic hosts within a population will be treated. Pyrogenic threshold–the parasite density at which a patient will exhibit fever as a symptom–varies widely between individuals, although increasing age and transmission intensity are both associated with higher pyrogenic thresholds [88, 89]. Here, hosts with merozoite density exceeding a threshold of 5000/*μ*L are considered to be symptomatic and thus become eligible for treatment. Symptomatic hosts are selected randomly to be treated for three days with a drug that kills 99.9% of sensitive parasites per day, which is sufficient to clear sensitive infections and is in line with expectations for artemisinin combination therapies [90]. Treatment persists even if the host recovers. Drug concentration is binary with no half-life.

### Simulations

#### Initialization and population burn-ins

The host population size is set at 100. Populations are initiated without vectors, mutation, or host mortality by infecting one host every five time steps for 250 time steps, producing a base population with a wide range of adaptive immunity and infection age. Parasite genomes begin with all bases set to A. After 250 time steps, host mortality is restored, with uniformly-distributed ages between 0 and 500, and mosquitoes are introduced, with uniformly-distributed ages between zero and 20. Given the other model assumptions, the mosquito age-structure will be retained throughout the simulations. The distribution of host ages, however, will shift over time due to background and infection-induced mortality. Populations are allowed to run for 2000 time steps. By the end of that time, measures of endemicity and strain diversity have reached equilibrium. At that time, drug treatment is introduced.

#### Emergent properties

The size of the mosquito population, set at either 300 or 1200 individual vectors, is the source of top-down regulation of transmission intensity. However, the realized transmission intensity is the product of interactions between treatment, frequency of resistance, immunity and genetic diversity, each of which is, in turn, affected by transmission intensity as well as interactions with one another. As a result, complex patterns can emerge from a few rules, and the nature of the interactions is difficult to intuit.

#### Resistance evolution dynamics

Because we model resistant parasites at both the within-host and the host population levels, we track the evolution of resistance by measuring prevalence of infections in the host population which are composed of at least 10% resistant parasites, a frequency which was sufficient to cause treatment failure in our simulations. Therefore, time to widespread treatment failure, given as Tf_a_;_l_, is defined as the time point at which 10% of infections are composed of at least 10% resistant parasites. Time to ubiquity, at which treatment failure predominates in a host population, is defined as the time point at which 75% of infections are composed of at least 10% resistance parasites. *T*_ubiq_ was only measured for conditions in which equilibrium prevalence of treatment failure exceeded 75%. In conditions under which equilibrium prevalence was below ubiquity, treatment failure transiently exceeded 75% in a handful of simulations; this was not counted. Simulations are run until they reach an equilibrium resistance prevalence.

#### Code and data availability

Code for the simulations was written in Python 3.6.9. Analysis was conducted using the R statistical package [91]. Treatment outcomes were compared using the Wilcoxon Rank Sum test, and normal approximated p-values are reported. For conditions in which the p-value was significant at the 95% confidence interval, treatment effect size was quantified using Vargha and Delaney’s A [92], implemented in the R library *effsize*. Simulation code is publicly available on Bitbucket and raw data is hosted at OSF.

## Supporting information

**S1 Fig Duration of untreated primary infections.**

**S2 Fig Parasite and immune dynamics during primary infections.**

**S3 Fig Infection trajectory during secondary exposure.**

**S4 Fig Strains exposed by age.**

**S5 Fig Complexity of infection by age.**

**S6 Fig The effect of recombination on time to resistance.**

**S7 Fig Evolutionary trajectory of costless resistance.**

**S8 Fig Frequency of resistance within each strain.**

## Acknowledgements

We thank Madeline Peters and Megan Greischar for helpful discussion and useful comments on the manuscript draft.

